# Geographic Distribution of a Missense Mutation in the *KRT38* Gene and Its Association with Heat Tolerance in Chinese Indigenous Cattle Breeds

**DOI:** 10.1101/2023.10.24.563762

**Authors:** Jialei Chen, Xin Liu, Jianyong Liu, Jicai Zhang, Bizhi Huang, Chuzhao Lei

## Abstract

**Context:** China has a vast area across many temperature zones and a variety of cattle breeds. These cattle resources are ideal models to research their adaptability to the environment. The *KRT38* gene is an acidic protein, and its coding product can be used as a component of hair production.

**Aims:** The objective of this study was to investigate the diversity of the *KRT38* gene in Chinese local cattle and the association of different genotypes with mean temperature (T), relative humidity (RH) and temperature humidity index (THI).

**Methods:** A missense mutation g.41650738 A > G in the *KRT38* gene was screened from the database of bovine genomic variation (BGVD), was genotyped in a total of 246 samples from 15 local cattle breeds in China by PCR amplification and sequencing. Finally, the correlation between the locus and the three climatic factors was analysed.

**Key results:** We successfully obtained the frequency of this SNP in three groups of cattle in northern, central and southern China. The frequency of allele A gradually declined from north to south, while the frequency of allele G showed the opposite trend with a clear geographic distribution.

**Conclusions:** Our results indicate that *KRT38* variation in Chinese indigenous cattle might be linked to heat tolerance.

**Implications:** Our analysis may support in finding out its importance as a genetic signal for heat tolerance in cattle reproduction and genetics.

## Introduction

Based on their geographical location and physical characteristics, the Chinese genetic resources for cattle can be divided into three main groups, which consist of 53 recognised native cattle breeds in the north, centre and south of China (Chen *et al*. 2018). In previous studies, indicine cattle are the descendants of the same ancestor as European taurine and African cattle, but they have experienced completely different evolutionary processes over the millennia (Verdugo *et al*. 2019). Natural selection has led to the evolution of heat resistance genes and stronger tolerance to warmer environments.

The global annual average temperature has risen gradually due to the the greenhouse effect, and the duration and intensity of high temperatures have also increased significantly. Temperature change can enhance heat stress on livestock, which results in a decrease in animal performance, such as the feed intake, milk production (West 2003) and reproductive performance (Hansen 2007). Animal heat tolerance is a quantitative trait (Gaughan *et al*. 2010; Li *et al*. 2011; Chang *et al*. 2012), and recent advancements in molecular genetic methodologies have facilitated the identification of the correlation between genetic variation at specific loci and their associated traits. Among these adaptive responses, the coat characteristics play a pivotal role in influencing the heat tolerance of animals under high ambient temperature and humidity conditions (Sarlo Davila *et al*. 2020).

Animal-derived filaments and hair fibers are composed of proteins, such as keratin and keratin-associated proteins (KAPs) (Gong *et al*. 2016), and also contain lipids and carbohydrates (Masukawa *et al*. 2005). *KRT38*, a gene belonging to the type II epithelial-keratin gene family, significantly influences the coat production of animals. Previous studies have revealed that several proteins in the keratin family affect the wool composition and structure (Yu *et al*. 2009; Li *et al*. 2018; Sulayman *et al*. 2018). In addition, keratins play a role in the formation of the hair shaft (Wu *et al*. 2008). Skin color and the thickness of the hair directly influence the thermotolerance of cattle that live in the tropics (Mattioli *et al*. 2000). *Bos indicus* has a smoother and shoter hair coat than *Bos taurus*. Due to these characteristics, *Bos indicus* regulates its body temperature and maintains cellular functions more efficiently during heat (Muchenje *et al*. 2008; Muchenje *et al*. 2009). *KRT38* is a potential candidate gene for wool traits (Sulayman *et al*. 2018), and genetic variation in the keratin and keratin-associated protein (KRATP) genes has been reported in many studies, indicating that the mutation contributes to phenotypic differences in wool (Langbein *et al*. 2007; Zeng *et al*. 2018; Zhang *et al*. 2018).

In contrast, there has been no research conducted regarding the correlation between temperature tolerance and genetic variation in the *KRT38* gene among indigenous Chinese cattle breeds. The purpose of this study was to evaluate the *KRT38* gene as a genetic marker for heat tolerance for cattle breeding and genetics, by investigating the diversity of the *KRT38* gene in Chinese native cattle and the relationship between various genotypes and mean temperature (T), relative humidity (RH) and temperature humidity index (THI).

## Materials and methods

### Ethical statement

The animal experimentation methods employed in this study were authorized by the Animal Care and Use Committee of the Institute of Animal Science, Northwest A&F University, Shaanxi, China, following the guidelines set forth in the field of biological sciences (Protocol number, WAFAC1008). During the sampling process, all samples collected in this study were examined with the consent of their owners and were standardized as far as possible.

### Animal Tissue DNA Extraction and Data Collection

A total of 246 ear tissues from 15 different native Chinese cattle breeds were collected from state-owned farms for this study (Table S1). Genomic DNA was extracted using the phenol-chloroform method (Xu *et al*. 2020). DNA samples were diluted to standard concentrations (50 ng μL^-1^) and stored at -80°C. Two environmental parameters (T and RH) over the last 30 years for the sampling sites of the 15 indigenous cattle breeds were collected from the Chinese Central Meteorological Office and were used to estimate heat tolerance traits (https://data.cma.cn/). The THI was calculated separately by the researchers based on the T and RH data. Based on nucleotide analysis of 15 indigenous cattle breeds, genotype and allele frequencies were determined. THI assesses the combined effects of T and RH to ascertain the heat load intensity of thermal climatic conditions, and it is a useful and simple measurement (Pinto *et al*. 2020). Next, it was calculated using the formula of National Oceanic and Atmospheric Administration:

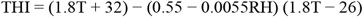

where T is the temperature in degrees Celsius and RH is the relative humidity as a percentage (Blanpain and Fuchs 2009).

Three climate variables with two genotypes each were subjected to environmental association studies, which used the general linear model (GLM) in SPSS 18.0. (Duricki *et al*. 2016). As Eckert suggested, we treated the environment variable as a phenotype. The statistical equation was:

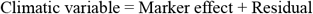

where climatic variables were the T, RH, and THI values between 1951 and 1980; marker effect was the fixed impact of the genotypes; and residual was the random element of the residual effect. Differences were considered significant at *P* < 0.01.

### Primer design, amplification, and novel SNP identification

The polymerase chain reaction (PCR) primer was designed based on the bovine *KRT38* sequence (GenBank accession rs210366642) by Primer Premier 5.0. The forward primer was 5’-AGGCTGGTCTCCAGTGTCAA-3’, and the reverse primer was 5’-CCTCAGTCCACCATGACTTCC-3’, which were designed to amplify a 114-bp product. The PCR amplification system consisted of 50 ng of genomic DNA, 10 μL of 2x PCR mix, 0.5 μM of each primer, and 8 μL of ddH_2_O (Jia *et al*. 2019). The following steps were involved in the cycling procedure: 5 min of 95°C denaturation; 35 cycles of 94°C for 30 s; 51°C for 30 s; and 72°C for 30 s primer extension; followed by an 8-min final extension at 72°C. The PCR products were identified directly at Shanghai Sangon Biotech Company, Shanghai, China, by electrophoresis on a 1%-agarose gel stained with ethidium bromide. SEQMAN TMIIv 6.1 was used to examine the sequencing outcomes.

### Statistical Analysis

The correlation between the three environmental factors (T, RH and THI) at the sampling sites was analyzed using SPSS software. This analysis employed a statistical linear regression model:

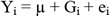

where Y_i_ is the value of T, RH and THI from 1950 to 1981; μ is the overall mean value; G_i_ = the fixed genotype effect; and e_i_ = the random residual effect. Differences were considered significant at *P* < 0.05.

Genotypic and allelic frequencies were estimated based on the observed genotypes in the examined breeds. Population indices such as homozygosity (Ho), heterozygosity (He), effective allele numbers (Ne) and polymorphism information content (PIC) were calculated. The THI, which considers the combined effects of T and RH, can serve as an indicator for assessing the intensity of heat load under thermal climatic conditions. This enables one to determine the heat load intensity more accurately.

Protein sequences and alignments for various species were obtained from the NCBI database (https://www.ncbi.nlm.nih.gov/). MEGA software (Kumar *et al*. 2018) was utilized to collect protein sequence similarities. Homology modelling of proteins before and after missense mutation was performed based on the bovine KRT38 protein sequence (NP_001070385.1) by the Amino Acid Explore website (www.ncbi.nlm.nih.gov).

## Results

### Novel missense mutation in the Chinese cattle *KRT38* gene discovered

In the current study, a novel A-to-G mutation (NC_037346.1 g.41650738 A > G) was identified in exon 1 of KRT38, resulting in the alteration of isoleucine at position p.I17T. This mutation was discovered in the Bovine Genome Variation Database and Selective Signatures (BGVD) available at (http://animal.nwsuaf.edu.cn/code/index.php/BosVar). Based on the sequence chromatograms, 26 Chinese cattle breeds exhibited three genotypes: AA (116), AG (72), and GG (51) (Fig. 1 and Table S1). In the present study, we assessed the genotypic and allelic frequencies of this gene in the cattle population (see to Table S2). The three genotypes exhibited relative frequencies of 0.4894, 0.3055, and 0.1850, respectively. The proportions of the A and G alleles were determined to be 0.7205 and 0.2795, respectively. A discontinuous distribution was also shown by the frequency of the allele G in *Bos taurus* and *Bos indicus*. From the geographical distribution of sampling sites, allele A in northern, central and southern Chinese cattle reached 0.9104, 0.6786 and 0.6031, respectively, and gradually diminished in Chinese indigenous cattle from south to north (Fig. 2).

**Fig. 1.**
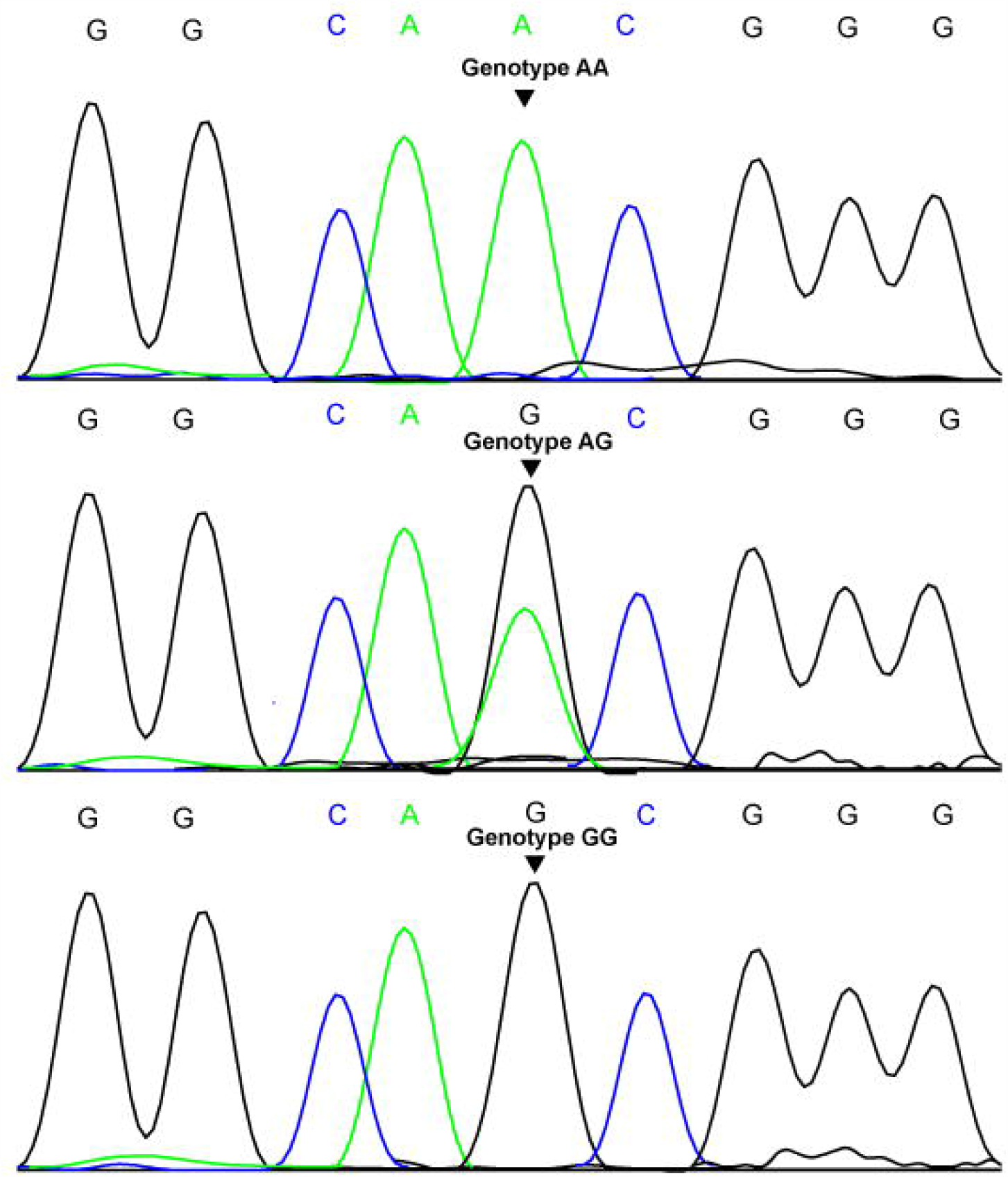
Identification of rs210366642 (NC_037346.1 g.41650738 A > G) mutations in the *KRT38* gene of bovines.

**Fig. 2.**
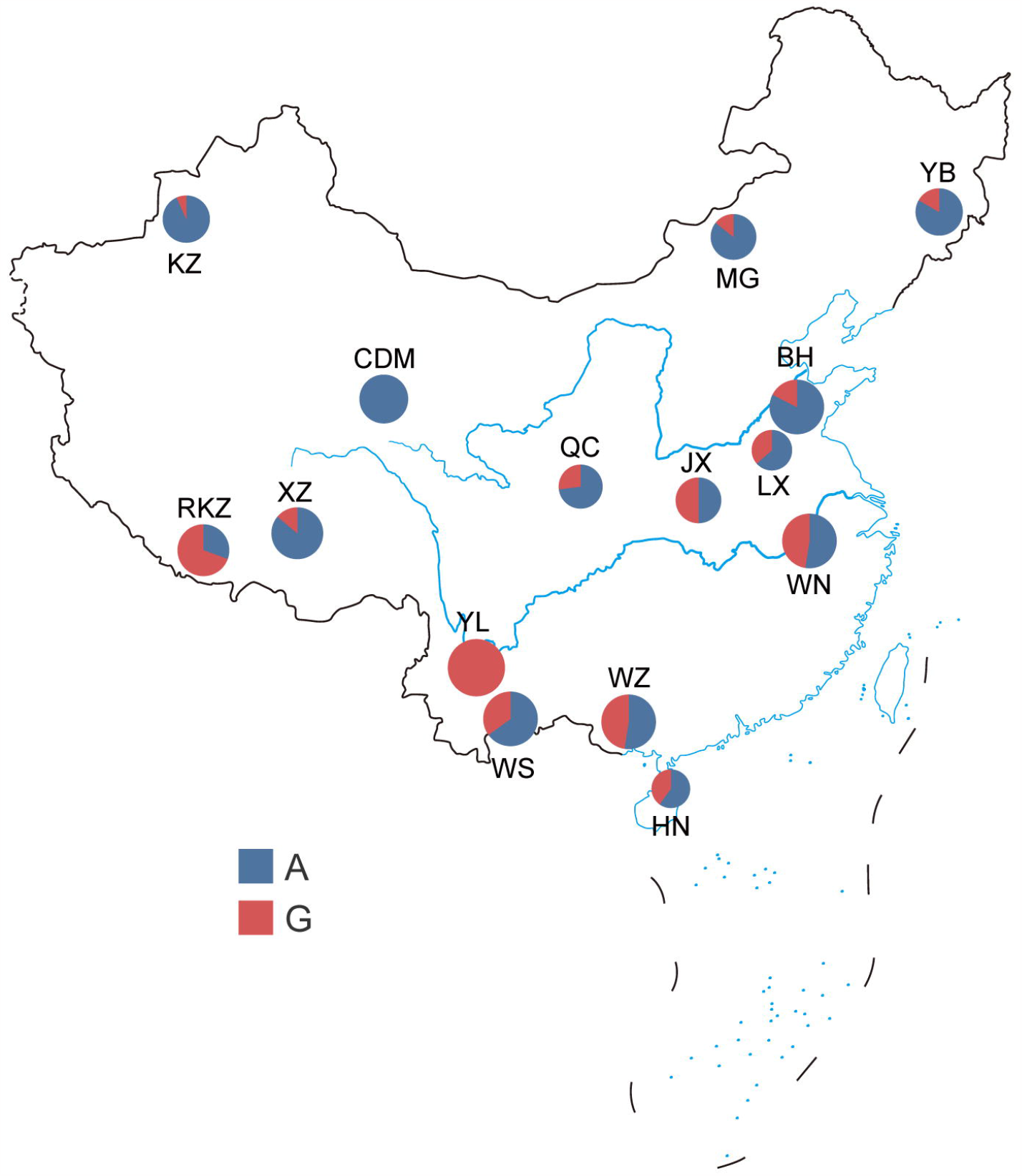
Geographical distribution of the A and G alleles of the locus rs210366642 (NC_037346.1 g.41650738 A > G) of the keratin 38 (*KRT38*) gene among 15 cattle breeds in China. Circled areas are proportional to the sample size. Blue and purple represent genotypes A and G, respectively. BH, Bohai Black; JX, Jiaxian red; LX, Luxi; MG, Mongolian; QC, Qinchuan; XZ, Tibetan; WN, Wannan; WS, Wenshan; YB, Yanbian; YL, Yunling; CDM, Chaidamu; HN, Hainan; WZ, Weizhou; RKZ, Rikaze; KZ, Kazakh.

### Genetic parameter analysis of the *KRT38* gene

The genetic indices of Ho, He, Ne, and PIC were calculated and presented in Table S3. Ho, which assesses genetic diversity and historical information of a population, ranged from 0.50 to 1, indicating increased heterozygosity. He values ranged between 0 and 0.50, while Ne values ranged between 1 and 2. The PIC values varied from zero to 0.375. The northern populations exhibited predominantly low or intermediate polymorphism, whereas the central herds and southern populations displayed mainly low or intermediate polymorphism at the locus, respectively, based on the PIC classification (PIC < 0.25, low polymorphism; 0.26 < PIC value < 0.5, intermediate polymorphism; and PIC value > 0.5, high polymorphism).

### Correlation Analysis of the *KRT38* Gene for Heat Tolerance

The correlation analysis results between the genotype g.41650738 A > G and three environmental parameters (T, RH and THI) were obtained from sample regions and genotypes of 246 Chinese cattle breeds. Table 1 presents the findings of this analysis. The genotypes AA and GG differed significantly (*P* < 0.01) from each other. Our data suggested that more individuals with the GG genotype than the AA genotype were present in hot, humid areas, indicating a relationship between allele G and heat tolerance in Chinese cattle. Based on the subject impact test on the *KRT38* genotypes (Table S4), RH and genotypes were found to be significantly related.

**Table 1.** Least squares mean and standard error for the temperature (T), relative humidity (RH) and temperature–humidity index (THI) of different genotypes of the single-nucleotide polymorphism (SNP) rs210366642 (NC_037346.1 g.41650738 A > G) in the *KRT38* gene.

| SNP | Genotype (n) | Temperature (°C) (LSM ± SE) | Relative Humidity (%) (LSM ± SE) | Temperature–Humidity Index (LSM ± SE) |
| --- | --- | --- | --- | --- |
| rs210366642:<br>(g.41650738<br>A > G) | AA (116) | 10.156±0.501<sup>A</sup> | 62.095<sup>A</sup>±1.353<sup>A</sup> | 52.492<sup>A</sup>±0.717<sup>A</sup> |
|  | AG (79) | 13.087±0.710<sup>B</sup> | 70.481<sup>B</sup>±1.119<sup>B</sup> | 56.513±1.053<sup>B</sup> |
|  | GG (51) | 13.230<sup>B</sup>±0.877<sup>B</sup> | 71.059<sup>B</sup>±1.255<sup>B</sup> | 56.689±1.267<sup>B</sup> |

LSM ± SE, the least square means with standard errors for diverse genotypes and environmental parameters. Means in the same column and locus with difference capital superscripts, A and B, are different at $P < 0.01$ .

### Mutation Analysis

Mammals possess complete coats that endure throughout their lifetimes and exhibit the ability to regulate their body temperature. Animals carrying the I17T mutation demonstrate enhanced heat tolerance compared to non-mutated animals, potentially attributable to functional modifications in *KRT38* caused by sequence variations. The alteration at Ile17 in the *KRT38* gene observed in cattle is identical to that found in zebu, yak, sheep and goats. Consequently, we proceeded to compare the protein sequences of KRT38 with those of other species (Fig. 3A). Our evolutionary analysis indicates that the I17T variation, identified in cattle, is rare among other mammalian species. Furthermore, we investigated the genetic pattern of this substitution (c.50 A > G, p.I17T) in cattle genomes from various geographical regions and observed a higher frequency of the mutation (c.50 A > G) in cattle residing in hot and warm climates, which aligns with the trend observed in China (Fig. 3B). The bovine KRT38 protein sequence was used to predict the protein three-dimensional structure by the homologous modelling method with the Amino Acid Explore website (www.ncbi.nlm.nih.gov). The missense mutation at g.41650738 A > G leads to a change in the amino acid encoding from isoleucine to threonine, which may lead to a change in protein properties (Fig. 4). Thus, we hypothesized that the substitution of residue 17 in the *KRT38* gene is probably related to heat tolerance, and this locus is likely to play a role in heat tolerance.

**Fig. 3.**
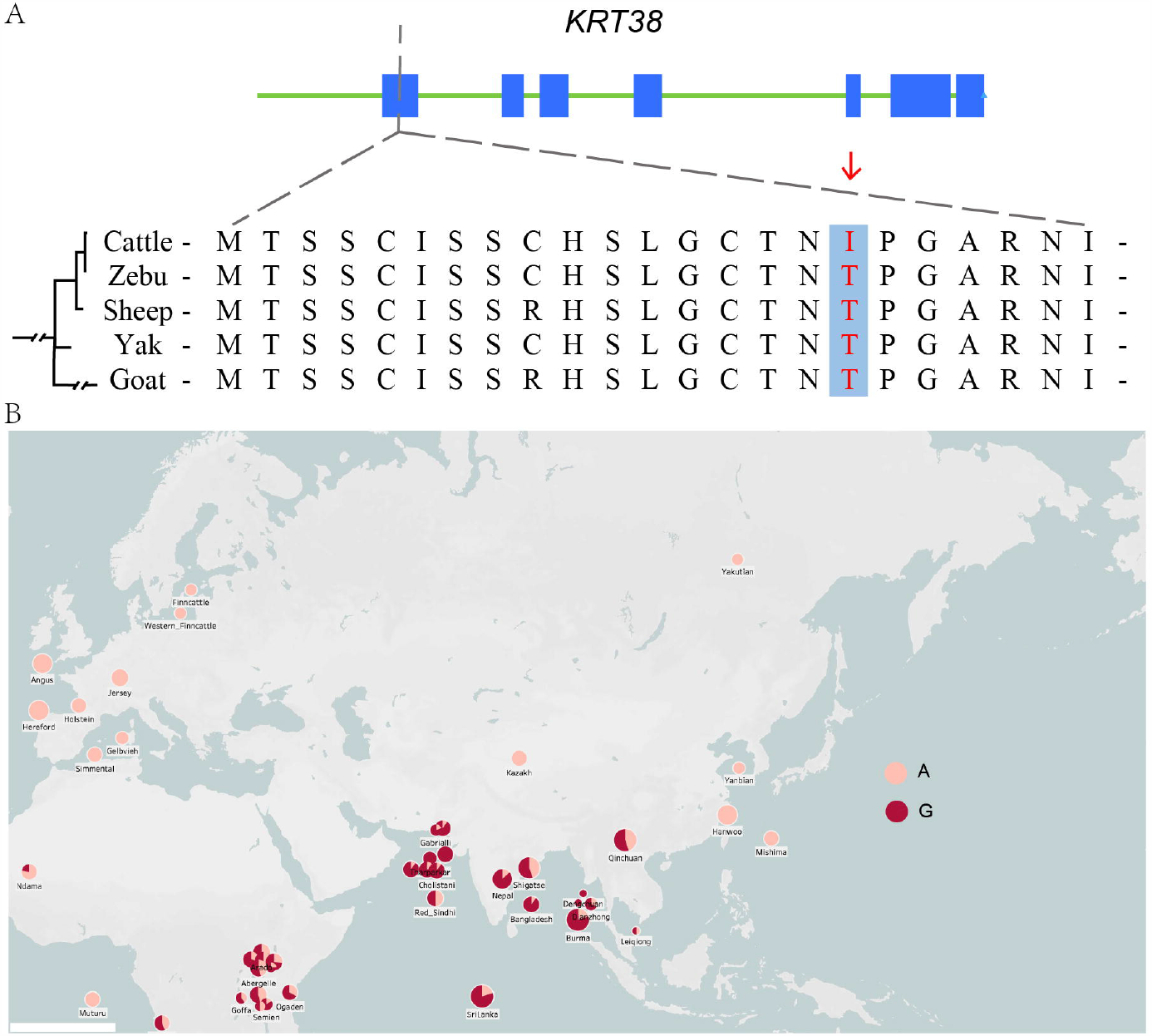
Genetic polymorphism of *KRT38* across five mammals. A: Graph depicting the bovine *KRT38* gene structure, showing the location of rs210366642 (NC_037346.1 g.41650738. A > G) mutation. Local alignment of exon 1 of the KRT38 protein showing the p. I17T mutation and adjacent amino acids in the five mammals. B: Genetic pattern of *KRT38* rs210366642 (NC_037346.1 g.41650738 A > G) in cattle genomes worldwide.

**Fig. 4.**
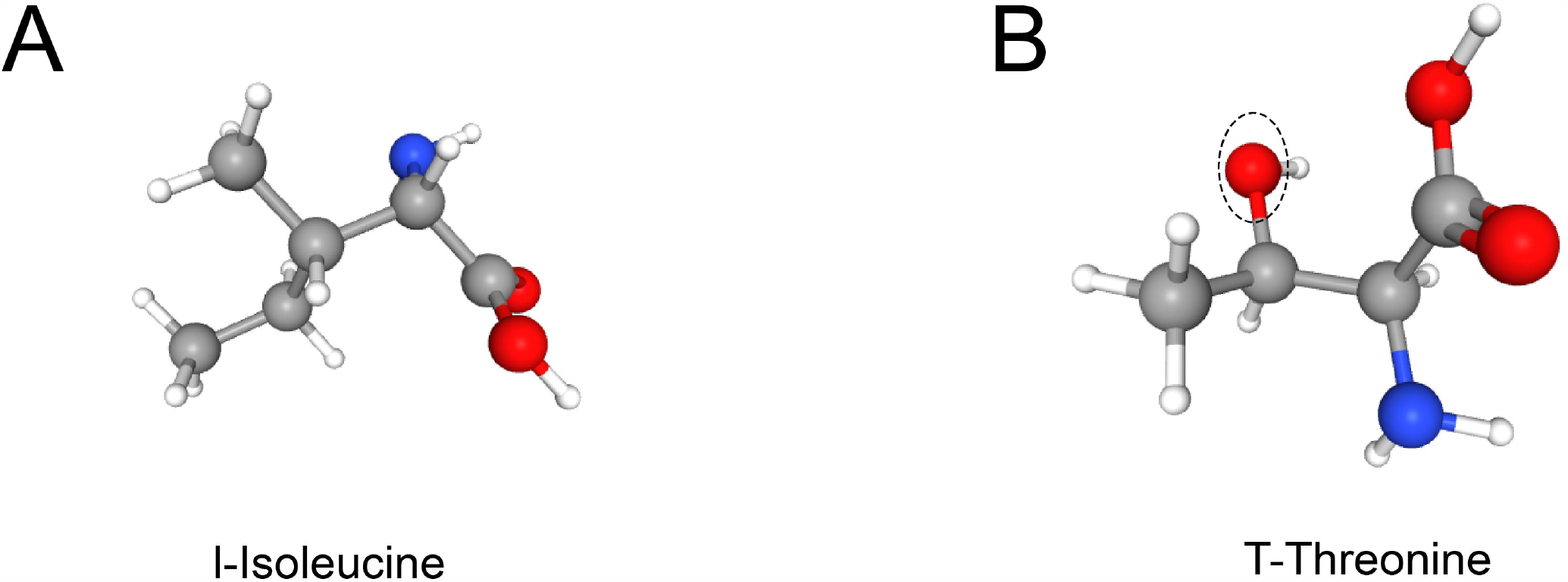
The tertiary structure prediction of KRT38 protein. A: wild type (Isoleucine) B: mutant type (Threonine).

## Discussion

Due to climate change and the imbalance between heat production and dissipation in some animals, the challenging tropical production environment could have determined more favorable bovine genotypes (Wolfenson and Roth 2019). Hair on mammalian skin serves multiple functions in thermogenesis, encompassing thermoregulation, sensory perception, and environmental defense (Blanpain and Fuchs 2009). A slick coat can be considered an indicative or indirect phenotype of various significant production qualities in tropical regions, as it is consistently associated with enhanced thermotolerance and higher milk yield in crossbreeds grazing under such conditions. This trait is regarded as a potential marker for the aforementioned production traits in tropical settings (Carabano *et al*. 2017). The performance of livestock under heat stress needs to be improved, and since thermal stress also affects animal productivity and genetic diversity among cow breeds negatively, it is essential to examine new methods and strategies. However, most economically important traits are quantitative by nature, meaning they are under the control of hundreds or thousands of genes (Meuwissen *et al*. 2001), with very few major genes having been identified as having an impact on these types of traits. In those cases, a genomic selection approach is the best method of genetic improvement. Identifying causal mutations for economically important traits would be beneficial for improving genomic selection, and hence, Meuwissen *et al*. (2022) have recently suggested new approaches to identify likely causal variants for complex traits by compensating for the very small effects with significantly larger sample sizes.

During the development of the epidermis, keratins play a crucial role as the predominant structural constituents (Wang *et al*. 2007). While a definitive correlation between the *KRT38* gene and heat stress effects in cattle is yet to be established, the site analysis and comparison of three environmental parameters (A and G) conducted in this study may imply a potential association between the mutation site of the *KRT38* gene and the heat tolerance of Chinese cattle. This observation aligns with the geographical distribution of hot and humid climates across China (Supplementary Table S2). The frequency of mutations at this SNP is higher in Indian cattle bloodlines than in Chinese taurine cattle bloodlines, according to the results of the association study. In Fig. 1, regions with more intense heat stress or greater light intensity had higher percentages of individuals with the GG or AG genotype, and these regions are also where the majority of the indigenous cattle with Indian cattle blood can be found. We speculate that this SNP is related to heat stress as the mutation site shows a correlation with temperature and humidity.

*Bos taurus* cattle are mainly found in northern China, while *Bos indicus* cattle are mainly found in southern China. *Bos taurus* and *Bos indicus* are both prevalent in central Chinese cattle due to trade and population mobility (Cai *et al*. 2007; Chen *et al*. 2018). While northern cattle breeds are often cold-tolerant, southern cattle breeds are typically resistant to thermal stress, as a prior study revealed. (Yudin *et al*. 2021). The examination of genotypic and allelic frequencies for the SNP (NC_037346.1 g.41650738 A > G) revealed that there was a significant difference in the regional distribution of the *KRT38* gene variation among the native Chinese cow breeds. In contrast to the A allele, which had a different pattern and corresponded with the distribution of indicine and taurine cattle in China, the frequency of the mutant G allele increased progressively from the southern region to northern China. This indicated how southern and northern breeds were affected by the introduction of the heat-resistant *Bos indicus* and the heat-sensitive *Bos taurus*, respectively. The unique geographical features of the Qinghai-Tibet Plateau have given rise to very distinctive breeds of cattle. In our study, we classified Tibetan cattle and Rikaze cattle from the Tibet region as special groups. Fig 2 shows that Tibetan cattle share most of their haplotypes with northern cattle. Interestingly, we found that the mutation G in Rikaze is consistent with the pattern of *Bos indicus*, which is consistent with previous research results that the Shigatse Humped have a pedigree of *Bos indicus* (Xia *et al*. 2019). The haplotype distribution of the special group indicates that the Qinghai-Tibet Plateau region has haplotype types of both *Bos taurus* and *Bos indicus*.

The GG genotype was more frequent in areas with higher T, RH and THI, as shown by the correlation between the novel SNP (NC_037346.1 g.41650738 A > G) and the three environmental parameters (T, RH, and THI). The GG genotype may be favored by the hot and humid climate in southern China, while the north may favor the AA genotype. Our research indicates that variations in bovine *KRT38* may have an effect on temperature tolerance. However, further testing is needed to testing different genotypes of animals and observe their physiological performance and response.

Furthermore, various studies have supported the notion that marked introgression between the bovine species yak, gayal, gaur, and banteng may help in adaptation to local conditions (Chen *et al*. 2018; Wu *et al*. 2018). China has been identified as a region where interbreeding between *Bos taurus* and *Bos indicus* has occurred, particularly in the central plains (Yue *et al*. 2014). In our research, the *KRT38* gene mutation stands as a unique exemplification of an amino acid residue convergent alteration that is shared by at least of 5 animal species. Thus, it is speculated that the cattle breeds in the central area carry allele G due to the occurrence of genetic mixing. The *KRT38* variant was the only cattle-specific missense mutation that was not observed in other closely related species or the Bovinae species in our study. We hypothesize that this genetic variation is exclusive to cattle. Our phylogenetic analysis reveals that the I17T mutation discovered in cattle is infrequent in northern China but has a higher likelihood of occurrence there. We propose that the KRT38 variation in cattle potentially exerts an influence on their skin pigmentation, hair length, and thickness, thus playing a pivotal role in enhancing their capacity to adapt to extreme ambient temperatures.

## Conclusions

Our findings have indicated that mutations within the *KRT38* gene have expanded the characterisation of genetic variations and their potential may be association with heat tolerance in indigenous Chinese cattle. However, it is important to note that the majority of adaptive traits related to environmental conditions are quantitative in nature and exhibit limited heritability. Furthermore, environmental adaptation is influenced by a multitude of genes. Therefore, it is crucial to continue identifying significant genes or loci that can enhance cattle productivity. These findings significantly contribute to our evolving comprehension of the adaptive genetic differences observed in cattle and other livestock species inhabiting diverse climatic regions.

## Data availability

Data availability is not applicable to this article, as no new data were created or analyzed in this study.

## Acknowledgements

We would like to thank the Yunnan Academy of Grassland and Animal Science, which contributed to the experimental design.

